# The role of a single non-coding nucleotide in the evolution of an epidemic African clade of *Salmonella*

**DOI:** 10.1101/175265

**Authors:** Disa L. Hammarlöf, Carsten Kröger, Siân V. Owen, Rocío Canals, Lizeth Lacharme Lora, Nicolas Wenner, Timothy J. Wells, Ian R. Henderson, Paul Wigley, Karsten Hokamp, Nicholas A. Feasey, Melita A. Gordon, Jay C. D. Hinton

## Abstract

*Salmonella enterica* serovar Typhimurium ST313 is a relatively newly emerged sequence type that is causing a devastating epidemic of bloodstream infections across sub-Saharan Africa. Analysis of hundreds of *Salmonella* genomes has revealed that ST313 is closely-related to the ST19 group of *S*. Typhimurium that cause gastroenteritis across the world. The core genomes of ST313 and ST19 vary by just 1000 single-nucleotide polymorphisms (SNPs). We hypothesised that the phenotypic differences that distinguish African *Salmonella* from ST19 are caused by certain SNPs that directly modulate the transcription of virulence genes.

Here we identified 3,597 transcriptional start sites (TSS) of the ST313 strain D23580, and searched for a gene expression signature linked to pathogenesis of *Salmonella*. We identified a SNP in the promoter of the *pgtE* gene that caused high expression of the PgtE virulence factor in African *S.* Typhimurium, increased the degradation of the factor B component of human complement, contributed to serum resistance and modulated virulence in the chicken infection model. The PgtE protease is known to mediate systemic infection in animal models. We propose that high levels of expression PgtE of by African *S*. Typhimurium ST313 promotes bacterial survival and bacterial dissemination during human infection.

Our finding of a functional role for an extra-genic SNP shows that approaches used to deduce the evolution of virulence in bacterial pathogens should include a focus on non-coding regions of the genome.

## Introduction

*Salmonella enterica* serovar Typhimurium (*S.* Typhimurium) is one of the best understood bacterial pathogens, and a major cause of gastroenteritis globally. One sequence type of *S.* Typhimurium, ST313, is the primary cause of invasive non-typhoidal Salmonellosis (iNTS) across Africa, resulting in ∼388,000 deaths each year^1^. Co-infection with HIV or malaria infection and young age (<5 years of age) are known risk factors for iNTS infection^1,2^.

Multi-drug resistance (MDR) has contributed to the expansion of *S.* Typhimurium ST313. Whole-genome sequence-based phylogenetics revealed clonal replacement of ST313 lineage 1 by lineage 2 in the mid-2000s, accompanied by the acquisition of chloramphenicol resistance^3^. The ST313 clade has recently acquired resistance to ceftriaxone, a first-line antibiotic for MDR bacterial infections^4^. Genomic comparison between the ‘classical’ gastroenteritis-associated *S.* Typhimurium ST19 and the African ST313 isolates shows that gene content and synteny are highly conserved, that ST313 has a distinct repertoire of plasmids and prophages, and carries 77 pseudogenes reflecting a degree of genome degradation^5,6^. ST313 and ST19 share >4,000 genes, and their core genomes differ by about 1,150 SNPs^4^. We have reported that 2.7% of the *S*. Typhimurium isolated from patients in England and Wales are ST313, but lack the characteristic prophages BTP1 and BTP5 that are signatures of African ST313 lineages^7^.

Certain virulence-associated phenotypes have been examined in ST313 strains. Compared with the ST19 group of gastroenteritis-associated *S.* Typhimurium, ST313 is more resistant to complement-mediated killing by human serum^8,9^ and to macrophage-mediated killing^10^. ST313 exhibits a stealth phenotype during macrophage infection consistent with an immune evasion strategy that causes reduced levels of IL-1β cytokine production, apoptosis and Caspase-1-dependent macrophage death^10,11^.

We used a functional genomic approach to search for single nucleotide polymorphisms responsible for the increased virulence of *S.* Typhimurium ST313 lineage 2.

## Results

The reference strain for *S.* Typhimurium ST313 lineage 2 is D23580, which was isolated from an HIV-negative Malawian child^5^. The strain 4/74 was isolated from a calf in the UK and is a well-characterised representative of *S.* Typhimurium ST19. Our challenge was to identify which, if any, of the >1000 SNPs that separate strains D23580 and 4/74 serve to differentiate the strains in terms of gene expression and phenotype. We investigated whether the emergence of the epidemic clade of *S.* Typhimurium ST313 was linked to the altered expression of a core genome-encoded virulence factor. Rather than focusing on a comparison of the core genome, we used comparative transcriptomics to identify transcripts that were both expressed at different levels and associated with a distinct SNP in the promoter region.

This study built upon the primary transcriptome of *S.* Typhimurium ST19 strain 4/74 which we determined using a combination of RNA-seq and differential RNA-seq (dRNA-seq) under multiple infection-relevant growth conditions^12,13^. By working at the single-nucleotide level, we defined transcriptional start sites (TSS), and catalogued the transcripts expressed in the bacterial cell^14,15^. Here, we used the same approach to define the primary transcriptome and to identify the transcriptional start sites (TSS) of D23580, a representative strain of African *S.* Typhimurium ST313. RNA was isolated from *in vitro* growth conditions that reflect the extracellular and intracellular stages of infection, namely early stationary phase (ESP) and the SPI2-inducing condition (InSPI2) (Methods). To find all relevant TSS, a pooled sample containing RNA from 16 environmental conditions was also analysed^13^ (Methods). TSS were identified by comparison of mapped sequence reads from each pair of dRNA-seq and RNA-seq samples as described^12,13,15^. We identified 3,597 TSS for *S.* Typhimurium strain D23580, revealing the active gene promoters across the genome of an ST313 isolate for the first time. Previously, we reported the locations of 3,838 TSS for the ST19 strain 4/74^12^. Categorisation of the TSS into different classes showed that a similar proportion of transcription initiation sites of 4/74 and D23580 were designated as primary (61%) or antisense (11%) (Figure 1A).

**Figure 1.**
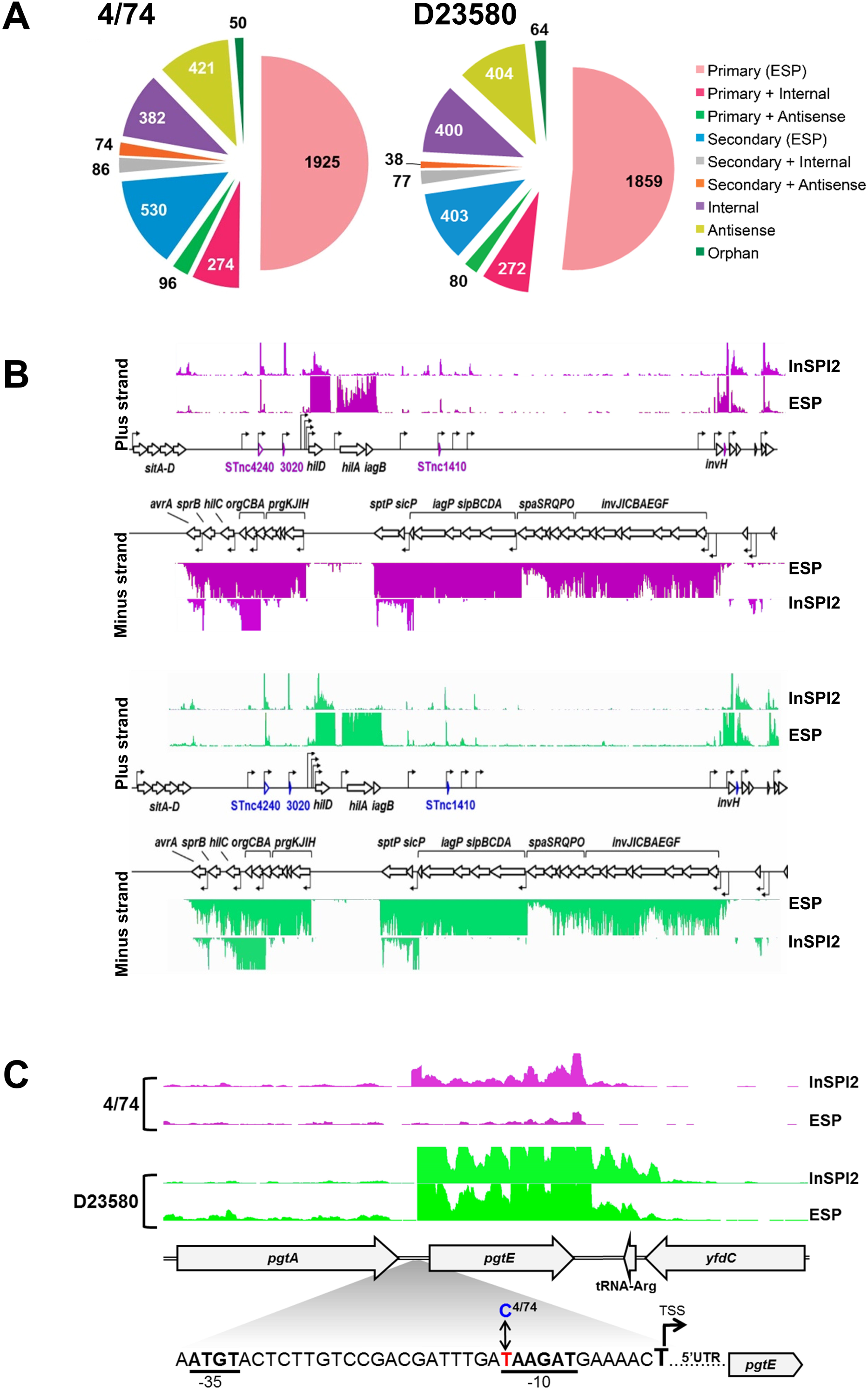
Primary transcriptome analysis of D23580 shows virulence gene *pgtE* is highly expressed, and is associated with a SNP in the conserved −10 promoter motif. Classification of Transcriptional Start Sites of *S*. Typhimurium in 4/74 and D23580. (A) Categorization of TSS identified in *S.* Typhimurium 4/74 and D23580, respectively, into nine different promoter classes^15^. (B) Visualization of Mapped Sequence Reads of the SPI1 Pathogenicity Island in *S.* Typhimurium 4/74 and D23580, respectively (IGB, scale 0–100 normalized reads for every sample). Names of coding genes and sRNAs are labelled in black and blue, respectively. TSS are indicated by arrows. (C) The sequence reads mapped to the *pgtE* locus were visualized in the Integrated Genome Browser^41^ (scale 0–100 normalized reads for every sample). Magnified region shows the pgtE promoter with −35/−10 promoter motifs in bold, and the T^D23580^ or C^4^/^74^ SNP highlighted.

We determined the level of conservation of transcriptional organisation between D23580 and 4/74 by identifying the TSS shared between the two strains. The locations of the majority of the TSS defined for strain 4/74 were conserved in strain D23580.

Specifically, of the 3,838 TSS of strain 4/74, 390 were absent from D23580 and included TSS located in the 4/74-specific regions of prophages sopEΦ and Gifsy-1 (Supplementary Table 1g). We identified 63 D23580-specific TSS, mainly located in the BTP1 and BTP5 prophages of D23580 which are absent from strain 4/74^5,6^ (Supplementary Table 1d).

To benchmark the transcriptional architecture, we first focused on *Salmonella* Pathogenicity Islands SPI1 and SPI2 which are required for key aspects of *Salmonella* virulence^16^. The locations of all TSS within the SPI1 and SPI2 islands were identical in strains D23580 and 4/74 (Figures 1B, Supplementary Table 1c). In summary, two closely-related *S.* Typhimurium strains that varied by ∼1,500 SNPs at the core genome level had a high level of conservation at the transcriptional level and shared 90% of promoter regions (Supplementary Table 1a).

To address our hypothesis that the level of expression of certain virulence genes varied between strains D23580 and 4/74 due to changes at the DNA sequence level, we cross-referenced the SNP differences between the two strains with the locations of the TSS. We identified 19 TSS which were associated with nucleotide polymorphisms in the −40 to −1 region of the 2,211 primary TSS of D23580 (Supplementary Table 3). We compared the expression level of each promoter between 4/74 and D23580, in 3 growth conditions, to identify the SNPs responsible for transcriptional changes. A SNP at the −12 position of the *pgtE* TSS, was associated with an average 11-fold increase in TSS expression in D23580 compared to 4/74 (Supplementary Table 3), and we investigated this experimentally.

### Identification of a nucleotide that modulates expression of the PgtE virulence factor

PgtE is an outer-membrane protease that belongs to the Omptin family^17^, cleaves and mediates resistance to alpha-helical antimicrobial peptides, and also disrupts the human complement cascade by degrading Complement Factor B and other proteins^18,19^. PgtE does not contribute to intra-macrophage replication *per se*, but stimulates bacterial dissemination during murine infection^20^ by facilitating extracellular survival upon release from host cells^21–24^. Expression of the *pgtE* transcript is induced during intra-macrophage replication^25,26^, controlled by the SPI2-associated regulators PhoPQ and SlyA ^18,27^, and is activated by OmpR/EnvZ and SsrA/B^14^.

The promoter and coding regions of the *pgtE* gene were compared between D23580 and 4/74 at the DNA sequence level, and differed by 2 SNPs. One SNP was identified in the coding region of *pgtE* at nucleotide location 2,530,498 in D23580 (2,504,548 in 4/74) generating a synonymous mutation [T54 (AC**T**) in 4/74→ T54 (AC**C**) in D23580]. The other SNP was located in the promoter region; the −12 nucleotide (relative to the +1 of the TSS) was C in 4/74 (C^4^/^74^) and T in D23580 (T^D23580^) (Figure 1C). This T nucleotide in the −10 motif is a highly conserved element of highly-expressed sigma70–dependent promoters^13^. We analysed the functional role of T^D23580^ in the *pgtE* promoter region experimentally by replacing the T^D23580^ nucleotide with C^4^/^74^ by single nucleotide exchange mutagenesis to generate strain D23580 *pgtE*^P4/74^. Whole genome sequencing confirmed that the D23580 *pgtE*^P4/74^ strain only contained the intended single nucleotide difference.

To determine the biological role of the T^D23580^ nucleotide, we assayed the level of *pgtE* transcription in 4/74, D23580, D23580 *pgtE*^P4/74^ and D23580 Δ*pgtE* strains using qRT-PCR (Figure 2A). The high level of *pgtE* expression in D23580 was reduced 10-fold by the introduction of the single C^4^/^74^ nucleotide in the −10 region of the *pgtE* promoter, P< 0.01 (Figure 2A). The level of the *pgtE* transcript expression in 4/74 and the D23580 *pgtE*^P4/74^ SNP mutant was similar.

**Figure 2.**
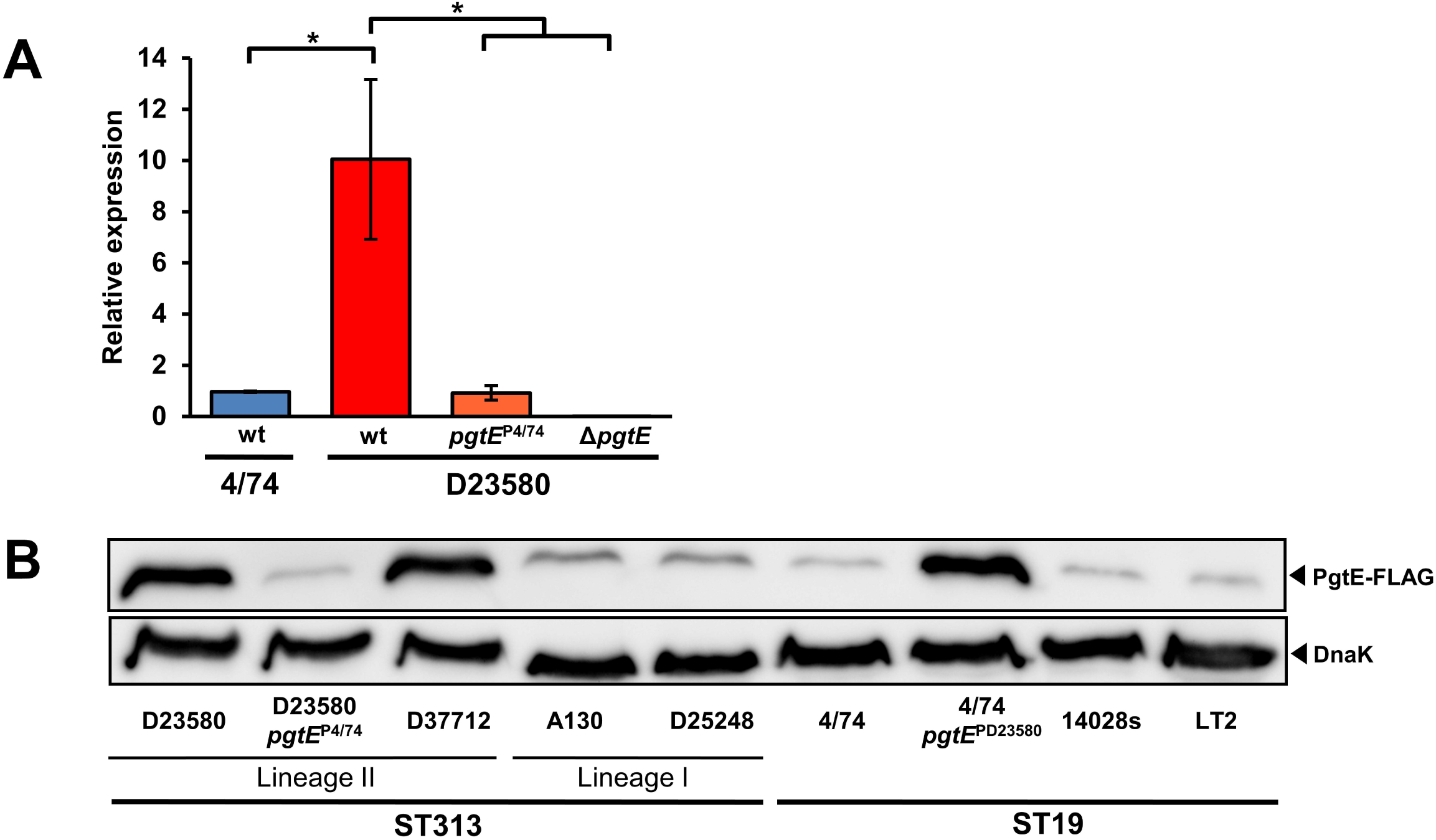
The T^D23580^ SNP in the *pgtE* promoter of *S.* Typhimurium is associated with increased *pgtE* transcription, PgtE protein production. (A) The level of *pgtE* transcript was measured by qRT-PCR and the relative gene expression, normalized to endogenous control *hns* was calculated using the *ddCt* algorithm^42^ and is the average of 3 biological experiments, with standard errors. Significant differences were analysed using an unpaired t-test (*p< 0.01). (B) Immunodetection by Western blotting of FLAG-tagged PgtE in representative strains of ST313 and ST19. The status of *pgtE* promoter (P^*pgtE*^) is only indicated for the strains with a mutated promoter. Detection of DnaK served as loading control.

We hypothesised that the high level of *pgtE* transcription would correlate with increased PgtE protein production in D23580 compared with wild-type strain 4/74 (Figure 2B). A second Lineage 2 isolate D37712 had the same PgtE phenotype. In contrast, low levels of PgtE were produced by the ST313 Lineage 1 isolates A130 and D25248, and ST19 isolates 14028 and LT2 (Figure 2B). The enhanced production of PgtE by D23580 was reduced to the level of the 4/74 strain by a single nucleotide change in strain D23580 *pgtE*^P4/74^. Consistent with this, the introduction of the T^D23580^ SNP into strain 4/74 caused increased PgtE protein production (Figure 2B). Taken together, our data show that D23580 expresses high levels of *pgtE* at the transcriptional and protein level, and this is driven by the T^D23580^ nucleotide in the −10 region of the *pgtE* promoter.

### The T^D23580^ SNP increases resistance to human serum killing and modulates cleavage of Complement Factor B

To determine the impact of the increased PgtE activity mediated by the promoter T^D23580^ SNP upon extracellular survival, we undertook serum bactericidal assays. Several bacterial factors contribute to the serum resistance phenotype of *Salmonella*, including the long heterogenic O-antigen side chains of smooth lipopolysaccharide (LPS), which is the outermost component of the cell envelope of the Gram-negative cell^28–30^. Therefore, we assayed resistance to human serum killing of *in vitro*-grown *S.* Typhimurium that lacked the LPS biosynthetic alpha1,3-glucosyltransferase enzyme WaaG. Following treatment with serum, the level of survival of D23580 *ΔwaaG* was significantly higher than D23580 *ΔwaaG pgtE*^P4/74^ (P=<0.05) (Figure 3A). No killing was observed following treatment with heat-inactivated serum that lacked active complement (data not shown). In summary, the promoter T^D23580^ SNP increases resistance of D23580 to serum killing and the low level of *pgtE* expression driven by the *pgtE*^P4/74^ promoter does not.

**Figure 3.**
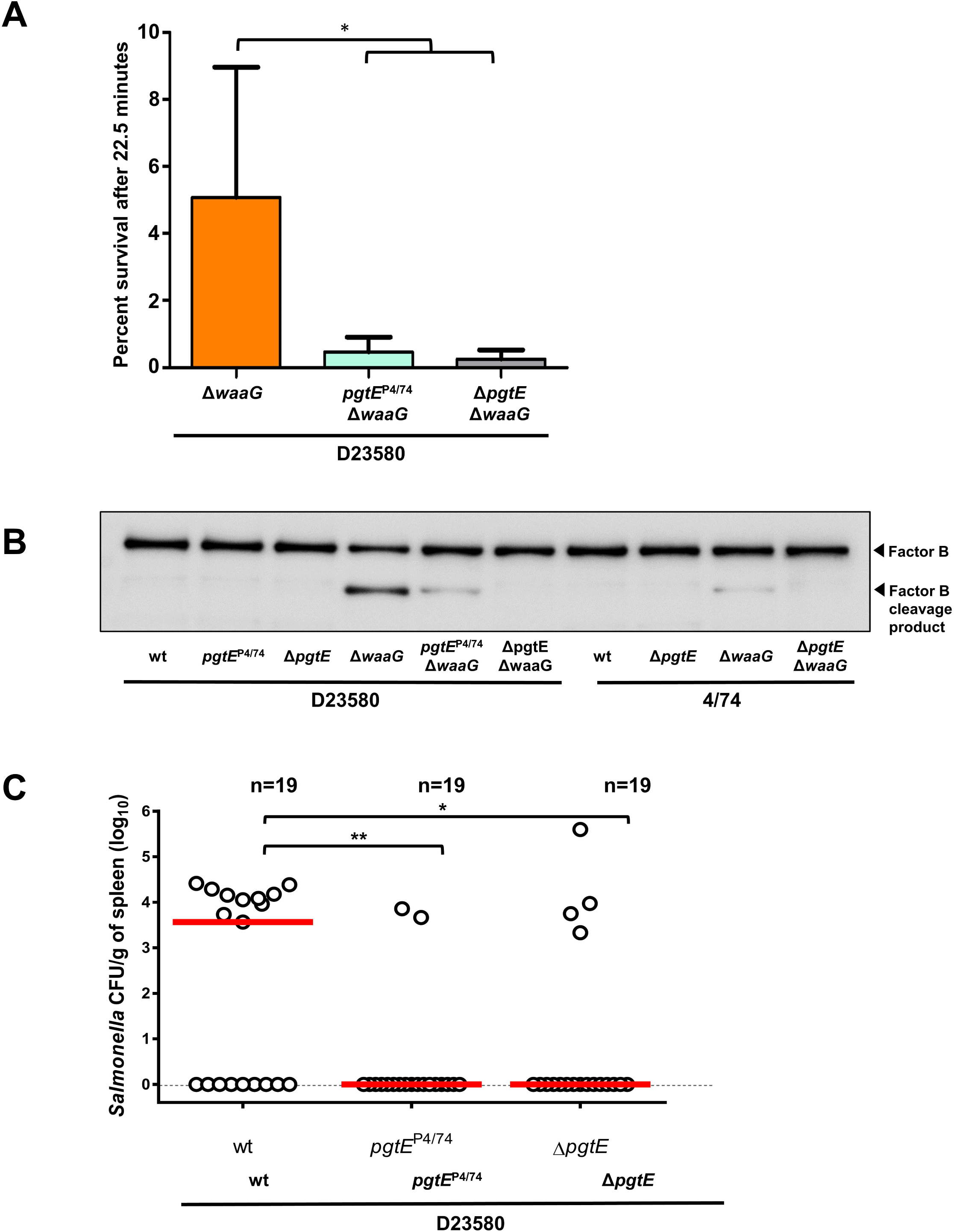
The *pgtE* promoter T^D23580^ SNP mediates increased resistance to human serum killing, enhances cleavage of human Complement Factor B and promotes virulence in the chicken infection model. (A) Sensitivity to pooled healthy human serum was assayed in a Δ*waaG* background (truncated LPS) to observe only the effect of outer membrane proteases. D23580 Δ*waaG* showed significantly less serum-sensitivity than D23580 *pgtE*^P4/74^ Δ*waaG* and D23580 Δ*pgtE* Δ*waaG* (P=0.04). (B) PgtE-dependent cleavage of Complement Factor B detected by Western blotting. Polyclonal antibody against factor B was used. (C) Viable counts of *Salmonella* Typhimurium D23580-derived strains as log CFU/g of spleen at 3 days post-oral infection (10^8^ CFU) of 7 day-old Lohmann Brown Layers. Data based on 19 individually sampled birds for each group; combined data for two separately repeated experiments. Each symbol represents the value for an individual chicken and the bars represent the median value for each group. Significance of differences between the groups was examined using a Mann–Whitney test. *, P<0.05; **, P<0.01.

To understand the mechanism of the serum-resistance phenotype, we determined the ability of the *S.* Typhimurium strains to mediate PgtE-dependent cleavage of Complement Factor B (Figure 3B). In agreement with the literature^31^, no PgtE activity was detected in strains expressing smooth LPS (Fig. 3C, Lanes 1-3 (D23580), Lanes 7-8 (4/74)). Because the results of the serum resistance assay showed that short (rough) LPS was required to visualise PgtE activity, we again conducted experiments in a *ΔwaaG* background (Figure 3B, Lanes 4-6 and 9-10), and determined that the D23580 *ΔwaaG* mutant showed a high level of Complement Factor B cleavage. In contrast, 4/74 *ΔwaaG* and the D23580 *ΔwaaG pgtE*^P4/74^ strains showed a low level of Complement Factor B degradation.

We speculate after the pathogen exits macrophages, the high level of expression of PgtE in *S.* Typhimurium ST313 strain D23580 interferes with opsonisation and increases resistance to complement-mediated serum killing.

### Assessment of PgtE-mediated virulence in the chicken infection model

Because *S.* Typhimurium ST313 has a hyper-invasive phenotype during chicken infection^32^, this infection model was used to assess the virulence of the wild-type D23580 and the D23580 *pgtE*^P4/74^ strains. Following oral infection, the D23580 *pgtE*^P4/74^ SNP strain and the D23580 *ΔpgtE* strain showed significant attenuation in comparison to D23580 wildtype (P=0.0035 and P=0.0379, respectively), based on two independent repeats of the experiment (Figure 3C). The data show some bird-to-bird variation between all three tested isolates, which is likely a consequence of the oral route of infection and the use of a commercial outbred chicken line. However, overall the results showed that exchange of the T^D23580^ SNP to the C^4^/^74^ genotype resulted in lower median bacterial numbers and a reduced number of animals with splenic infection, equivalent to that seen in the absence of PgtE (10/19 for D23580 compared to 2/19 for D23580 *pgtE*^P4/74^ and 4/19 for D23580 *ΔpgtE*). We conclude that PgtE is required for successful infection of the chicken, and that full virulence of D23580 requires the high levels of expression of PgtE driven by the T^D23580^ nucleotide.

### The *pgtE* promoter SNP is only carried by African ST313 lineage 2, and not lineage 1

To determine if the *pgtE* promoter T^D23580^ SNP is a characteristic feature of iNTS, the *pgtE* promoter SNP was analysed in the context of a phylogeny of 268 genomes of *S.* Typhimurium ST313 including isolates from Malawi, as well as recently described UK-ST313 genomes^7^. The 228 genomes that carried the T^D23580^ SNP formed a monophyletic cluster that included lineage 2, as well as the UK-ST313 strains that share most recent common ancestry with lineage 2 (Figure 4). The C^4^/^74^ SNP was found to be conserved in all 27 lineage 1 genomes and the UK-ST313 genomes which shared more recent common ancestry with lineage 1. This suggests that the T^D23580^ SNP first arose in a common ancestor of lineage 2 and a subset of the UK-ST313 (Supplementary Figure 2).

**Figure 4.**
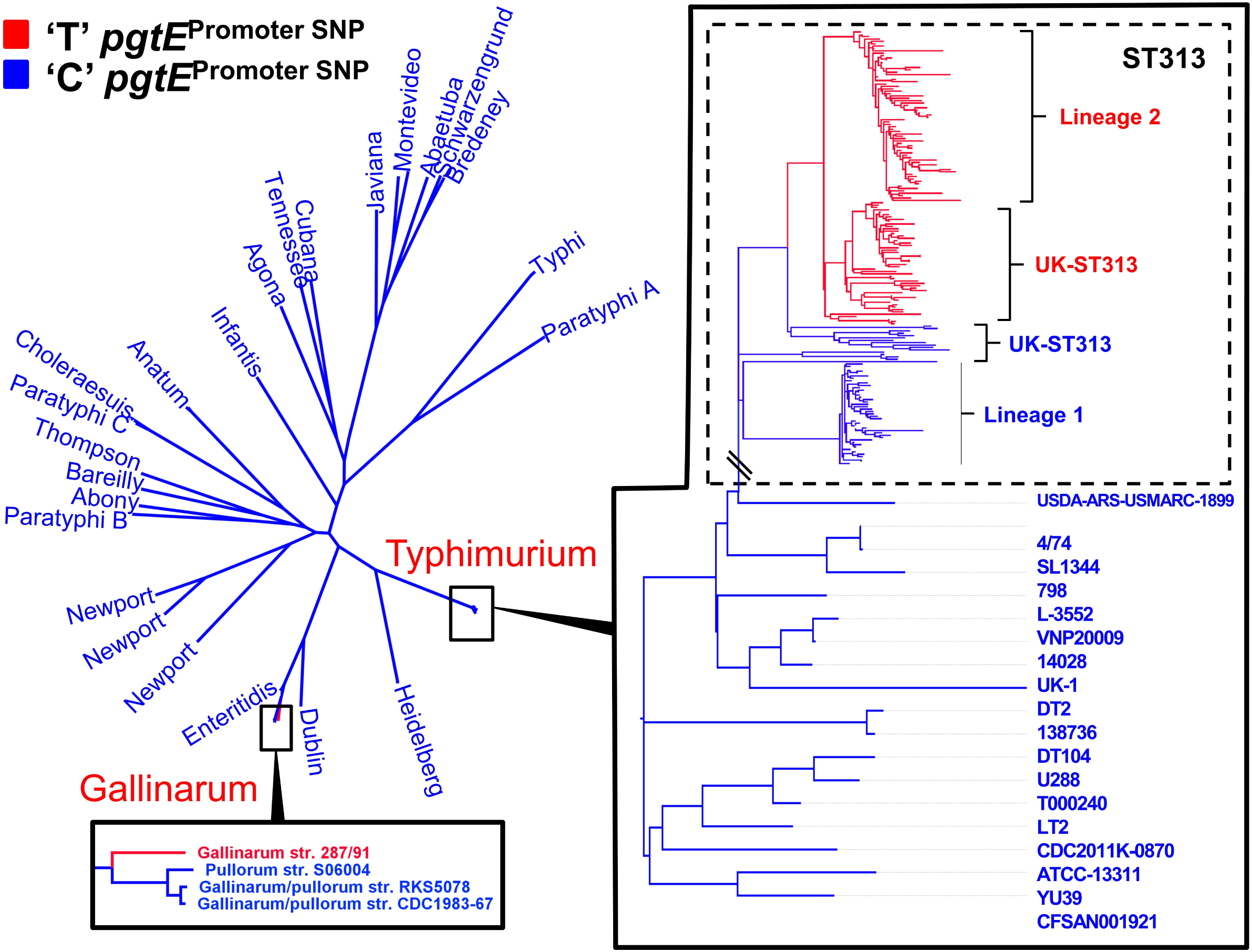
Conservation of the *pgtE* promoter −10 T^D23580^ nucleotide across *S. enterica* ssp. enterica. Maximum likelihood phylogenetic tree of 266 *S. enterica* genomes. Presence of C or T nucleotide at the −10 position of the *pgtE* promoter is indicated by blue and red respectively. The T nucleotide is found in 100 % of ST313 lineage 2 genomes surveyed. Outside of serovar Typhimurium, the T nucleotide is also present in the genome of Gallinarum isolate 287/91.

### The *pgtE* promoter is highly conserved in *Salmonella enterica*

To understand the wider distribution of the *pgtE* promoter SNP in the *Salmonella* genus, the conservation of the SNP was assessed in 84 published complete genomes representing the known genomic diversity of *Salmonella*. The *pgtE* TSS was not found to be conserved in *S.* Bongori (Supplementary Table 4). Of 80 *S. enterica* genomes screened, 79/80 genomes carried the C^4^/^74^ genotype. The T^D23580^ SNP was only found in *S.* Gallinarum str. 287/91, raising the possibility that the SNP has arisen independently in this serovar (Figure 4). Apart from the −12 SNP present in *S.* Typhimurium ST313 and *S.* Gallinarum, the *pgtE* TSS −35 region was found to be 100% conserved in 75/80 *Salmonella enterica* genomes, with only serovar Agona and subspecies Arizonae showing sequence divergence (Supplementary Table 4).

In summary, the −10 C^4^/^74^ → T^D23580^ allele of the D23580 *pgtE* promoter causes an increase in transcription of the *pgtE* gene and the production of high levels of PgtE protein in D23580. The increased activity of PgtE in D23580 leads to degradation of Complement Factor B which is required for activation of the alternative complement pathway. Importantly, the single SNP in the D23580 *pgtE* promoter drives the ability of D23580 to cause hyper-invasion in an avian infection model. Only one of 80 complete *S. enterica* genomes, that of *S.* Gallinarum, carried the −12 T allele. *S.* Gallinarum is the causal agent of fowl typhoid, suggesting a putative link between the T^D23580^SNP and the ability of *S*. *enterica* to cause systemic infection^33^. We have shown that the *pgtE* promoter SNP is a signature of ST313 lineage 2, which clonally-replaced ST313 lineage 1 in the early 2000s^3^. All isolates of ST313 lineage 2 carried the same −12 T^D23580^ SNP.

## Discussion

Previous studies have identified SNP mutations associated with the host tropism of notorious pathogens such as *Staphylococcus aureus*^34^, and *Campylobacter jejuni*^35^. However, these examples involved SNP mutations located within coding genes, and functionally-important SNP mutations have rarely been identified in intergenic regions of bacteria. As the expression of a gene is dependent on the −10 and −35 recognition motifs of sigma 70-dependent promoters^36^, a single nucleotide change can modulate promoter function. Examples include the C → T transition in the −10 promoter motif of the *Mycobacterium tuberculosis eis* gene, that increases *eis* expression to generate low-level resistance to Kanamycin^37^. Similarly, a promoter SNP that affected expression of *E. coli* succinate transporter *dctA* evolved to increase the utilisation of citrate as a carbon source in one population of the Long Term Evolution Experiment^38^.

Here, we have identified a single SNP responsible for high levels of expression of the PgtE outer membrane protease, and have linked this to the virulence of African *S.* Typhimurium ST313. Our study has implications for other bacterial genome-wide association studies, which should clearly include a focus on non-coding regions of the genome. The findings also emphasise the value of identifying all gene promoters in bacterial pathogens, to allow nucleotide differences to be correlated with the process of transcriptional initiation.

We propose that the high level of PgtE activity in D23580, together with the inactivated *sseI* effector gene^39^ and the acquisition of chloramphenicol resistance^3^, has been a factor in the success of epidemic ST313 lineage 2.

A pre-requisite for PgtE activity is the remodelling of LPS that occurs during intra-macrophage replication that results in shortening of the oligosaccharide chains^21^. *S.* Typhimurium bacteria produce high levels of *pgtE* transcript inside host macrophages^25,26^, and PgtE protease activity is high in bacteria released from infected macrophages^21^. Therefore the *pgtE* T^D23580^ SNP represents a putative mechanism for priming intracellular bacteria for an extracellular lifestyle, and the survival of complement-mediated attack by the innate immune system. The opsonic activity of complement has been shown to be essential for phagocyte-mediated killing of *Salmonella* in the blood of African people^40^ and therefore our data are consistent with the hypothesis that subversion of complement activity contributes to the pathogenesis of invasive non-typhoidal *Salmonella* in Africa.

The T^D23580^ SNP in the −10 motif of the *pgtE* promoter causes increased PgtE protease activity, and was an early evolutionary event in an ancestor of ST313 lineage 2 which primed the emergence and dominance of ST313 lineage 2 in iNTS disease across sub Saharan Africa.

## Material and methods

### Bacterial strains, growth conditions

*Salmonella enterica* serovar Typhimurium (*S.* Typhimurium) strain 4/74 (accession number CP002487), a representative of non-typhoidal *Salmonella* sequence type 19, and D23580 (accession number FN424405), a representative strain of non-typhoidal *Salmonella* sequencing type 313 (ST313) were used in the study. Strain D23580 was isolated from an HIV-negative child from Malawi with blood stream infection, and use of this strain has been approved by the Malawian College of Medicine (COMREC ethics number P.08/14/1614). Other wild-type strains belonging to ST19 and ST313 used in this study are listed in Supplementary Table 5.

All environmental growth conditions were repeated exactly as previously described^12^ with the exception that the ‘pool’ sample was obtained by pooling RNA from 16 environmental conditions (EEP, MEP, LEP, ESP, LSP, 25°C, NaCl shock, Bile shock, Low Fe^2+^ shock, Anaerobic shock, Anaerobic growth, Oxygen shock, NonSPI2, InSPI2, Peroxide shock (InSPI2) and Nitric oxide shock (InSPI2)^12^.

When required, Lennox broth (LB) was supplemented with the following antibiotics: chloramphenicol (Cm), 25 μg/ml; kanamycin (Km), 50 μg/ml, tetracycline (Tc), 20μg/ml and gentamicin (Gm), 20 μg/ml.

### Preparation of cDNA libraries and Illumina sequencing

Prior to RNA-seq, total RNA was extracted using Trizol, and treated with DNase I as described previously^13^. RNA integrity was inspected visually with the Bioanalyzer (Agilent technologies). Contaminating DNA was removed using DNase I (Ambion) and RNA samples were not ribo-depleted prior to cDNA library construction. cDNA library construction, TEX-treatment for dRNA-seq (ESP, InSPI2 and pooled sample) and RNA-seq with the Illumina HiSeq platform was carried out by Vertis Biotechnologie, Germany. All protocols were identical to those used previously^12,13^. Sequence reads were mapped against the *S*. Typhimurium D23580 reference genome using Segemehl, with accuracy set to 100%^43,44^. RNA-seq and dRNA-seq data can be downloaded as raw reads (.fastq file format) the GEO database accession number XXXXXXXX.

### Identification of Transcriptional Start Sites by a combination of RNA-seq and dRNA-seq

Methods used to assign TSS in D23580 have been described previously^13^. Briefly, a TSS was assigned when it was enriched in one of the dRNA-seq libraries (ESP, InSPI2 or Pool) compared with the corresponding RNA-seq library, and was linked to an expressed transcript. This analysis was followed by a second step of validation, in which the Transcripts Per Million (TPM) approach^45,46^ was used to calculate an expression value for the first 10 nucleotides associated with each TSS, designated the Promoter Usage Value (PUV)^13,26^. A TSS was considered to be expressed when the PUV was ≥10. A TSS was defined as ‘conserved’ between D23580 and 4/74 if the TSS nucleotide sequence was present in both strains, and the PUV value of the TSS was ≥10.

### Identification of TSS-located SNP mutations associated with different levels of transcript expression in D23580 and 4/74

A list of SNP differences between the D23580 and 4/74 reference genomes (accessions FN424405 and CP002487) was generated using NUCmer^47^ resulting in 1,488 SNPs and small indels (Supplementary Table 2). We identified the SNPs located within the −40 to −1 region of primary TSSs in D23580 (primary = primary + primary/as + primary/internal) (Supplementary Table 1, Supplementary Table 3). PUV values for each promoter in D23580 and 4/74^13^ were used to analyse the activity of the promoters associated with SNP differences in the −40 to −1 TSS region.

### Bacterial strain construction using λ *red* recombineering

All the bacterial strains and plasmids used and constructed in this study are described in Supplementary Table 5 and the ssDNA oligonucleotides (primers) in Supplementary Table 6.

The Δ*pgtE*, Δ*waaG* and *pgtE-FLAG* mutations were constructed in *S*. Typhimurium ST19 and ST313 strains using the standard λ *red* recombination methodology^48^. The heat-inducible λ *red* recombineering plasmid pSIM5-*tet* was used and the induction of the λ *red* operon was achieved by heat treatment (42°C, 15 min) of bacterial cultures grown to mid-exponential phase (OD_600_ 0.3-0.4 at 30°C) in LB supplemented with Tc^48–50^. After recombination^48–50^, all the genetic constructs (except the Δ*waaG*::Kan mutation) were transferred into a clean wild-type background by phage transduction, using the P22 HT 105/1 *int-*201^51^ as previously described^6^. When required, the antibiotic resistance cassettes were flipped-out using the FLP recombinase expression plasmid pCP20-TcR^52^.

The *pgtE* gene was deleted in *S.* Typhimurium using PCR fragments generated with the primers DH95 and DH96 and the template plasmids pKD4 and pKD3. The resulting fragments, carrying respectively the Km resistance (Kan) or the Cm resistance (Cam) cassettes were respectively electroporated into D23580 and 4/74 carrying pSIM5*-tet* and recombinant *ΔpgtE*::Kan/Cam mutants were selected on Km or Cm LB agar plates. Finally, the mutations were transduced in the corresponding wild-type strain and the resistance cassette was removed, as described above. Similarly, *waaG* was inactivated using the primers del_waaG_F and del_waaG_R and pKD4 as template.

The FLAG-tagged strains were generated using a forward primer (DH93) which included the region homologous to *pgtE* end (‘*pgtE*), the nucleotide sequence encoding the FLAG octa-peptide in frame with the *pgtE* coding region, the *pgtE* stop codon and a region homologous to the resistance cassette (Song, Kong et al. 2008). The ‘*pgtE-*FLAG-Kan and ‘*pgtE-*FLAG-Cam modules were amplified by PCR using respectively pKD4 and pKD3 and primers DH93 / DH94. The resulting amplicons were respectively electroporated into *S.* Typhimurium ST313 (except A130) or ST19 (and A130) strains, carrying all pSIM5-*tet* and recombinants were selected on Km or Cm LB agar plates. The insertions were then transduced in to the corresponding wild-type strains and the resistance cassettes were removed, as described above.

### Construction of scarless single nucleotide substitution mutants

Two different methods were used to construct single nucleotide mutants. The single nucleotide T→C substitution in the *pgtE* promoter of *S.* Typhimurium D23580 was constructed using a single stranded DNA (ssDNA) recombineering approach, as has been previously described ^53^. The protocol was identical to that used for construction of mutants using λ *red* recombineering (described above) except that 400 ng of the manufactured primer DH90), (HPLC purified) were used in the transformation reaction. After 2 hours of recovery at 30°C, dilutions of transformation were plated on LB agar (without selection). Clones were re-streaked and screened by a stringent PCR with primers DH40 and DH41. Primer DH40 has full complementarity with the sought after mutant, representing one mis-match to the original strain (these two types could be distinguished using a stringent annealing temperature) while DH41 has full complementarity with both types of clone. The correct allele of the D23580 *pgtE*^P4/74^ strain was confirmed by whole genome sequencing using Illumina technology (MicrobesNG, University of Birmingham). Variant-calling analysis confirmed that the D23580 *pgtE*^P4/74^ strain had the intended single nucleotide difference compared with the WT strain (data not shown).

The single nucleotide C→T substitution in the *pgtE* promoter of *S.* Typhimurium 4/74 (chromosomal position 2504765) was carried out by a scarless genome editing technique based on the pEMG suicide plasmid, as previously described ^6,54^. The pEMG derivative pNAW41 that carries the *pgtE* promoter region with the specific substitution was constructed as follows: the regions flanking the targeted nucleotide were PCR amplified with the primers pairs NW_122 / NW_123 and NW_124 / NW_125, using 4/74 genomic DNA as template. The primers NW_123 and NW_124 encode for C→T substitution and are complementary to each other over a stretch of twenty nucleotides. The resulting PCR fragments (504 and 505 bp, respectively) were fused by overlap extension PCR and the resulting 989 bp fragment was digested and cloned into pEMG using the *Bam*HI and *Eco*RI restriction sites. The pNAW41 suicide plasmid was mobilized from *E. coli* S17-1 λ*pir* into *S.* Typhimurium 4/74 by conjugation and transconjugants that have integrated pNAW41 by homologous recombination were selected on minimal medium M9 agar supplemented with 0.2 % of glucose and Km. The resulting merodiploids were resolved using the pSW-2 plasmid as previously described^6^ and the C→T substitution was confirmed by PCR amplification and sequencing, using the primers NW_155 and NW_156.

### Quantitative PCR (qRT-PCR)

Total RNA was extracted from bacteria from mid-exponential (OD_600_ = 0.3) cultures of bacteria grown in PCN/InSPI2, DNase-treated with Turbo DNA-free kit (Ambion) and the RNA integrity was inspected visually using the Bioanalyzer. Complete DNA-removal was confirmed by a negative PCR reaction with 40 cycles. Four hundred ng of RNA was converted into cDNA using the GoScript Reverse Transcription System (Promega), with random primers according to the manufacturer’s instructions. The Sensifast SYBR Hi_ROX Kit (Bioline) was used for qRT-PCR. For each qRT-PCR reaction, performed in duplicate, 26.66 ng cDNA was used in total reaction volumes of 20 μl. The amount of *pgtE* and *hns* mRNA was calculated using a standard curve based on ten-fold dilutions of genomic DNA (10 ng/μl - 0.0001 ng/μl), included in each qRT-PCR run. The amount of *pgtE* mRNA was normalized to the amount of *hns* mRNA.

### Western blot analysis

*Salmonella* strains carrying the FLAG-tagged version of *pgtE* were grown in PCN/InSPI2 medium to OD_600_ = 0.3. Bacteria were harvested from 10 ml of culture by centrifugation (7,000 x *g*, 5 min, 4°C). Cells were washed once with PBS and suspended in 67.5 μl of the same buffer. Seventy-five microliters of Laemmli Buffer 2× (120 mM Tris-HCl pH 6.8, 4% w/v SDS, 20 % v/v glycerol, Bromophenol blue 0.02 % w/v) and 7.5 μl β-mercaptoethanol were subsequently added to the samples. The extracts were boiled for 10 min, chilled on ice, and cell debris were pelleted by centrifugation (20,000 × *g*, 5 min, 4°C). Fifteen microliters of the samples (supernatant) were loaded on a SDS 10% polyacrylamide gel and proteins were separated for 80 min at 150 V in SDS-PAGE running buffer (25 mM Tris, 192 mM glycine, 0.1% SDS). Proteins were transferred onto a methanol soaked PVDF membrane (Roche, Cat. No. 3 010 040) using semi-wet transfer system (Bio-Rad, #170-3940) for 2 hours, 125 mA at 4°C in transfer buffer (25 mM Tris, 192 mM glycine). The membrane was blocked for 15 hours at 4°C in Tris-buffered saline (TBS: 10 mM Tris-HCl pH 7.5, 0.9% NaCl) supplemented with 5 % w/v of dry skimmed milk. Incubation with the antibodies (1 hour at room temperature) and washing steps (two washes for 15 min each) were done in TBS containing 0.1 % v/v Tween 20 and 0.5% w/v dry skimmed milk. The primary antibodies Monoclonal ANTI-FLAG M2 antibody (1:3,000 diluted, Sigma-Aldrich F3165) and anti-DnaK mAb 8E2/2 (1:10,000 diluted, Enzo Life Sciences, ADI-SPA-880) were used for the detection of PgtE-FLAG and DnaK (loading control), respectively. After incubation with the primary antibodies, the membrane was washed and incubated with the secondary antibody (Goat anti-mouse IgG (H+L)-HRP, 1:2,500 diluted, Bio-Rad, # 172-1011). After washes, the membrane was rinsed briefly in TBS prior to addition of Pierce ECL western blotting substrate (Thermo Scientific, 32109) and the chemiluminescence reaction was measured using the Image Quant LAS 4000 imager (GE Healthcare Life Sciences).

### Complement Factor B cleavage assay

The Complement Factor B cleavage assay was carried out as described^19^, with the following modifications. Bacterial strains were grown overnight in LB at 37°C. Overnight culture was pelleted by centrifugation and washed twice in PBS. Two OD_600_ (in 200 μl PBS containing 33 ng/μl of Complement Factor B) were incubated at 37°C in a heat block with agitation (700 rpm) for two hours and subsequently pelleted by centrifugation. Sixty-six ng of Complement Factor B were separated on an SDS-PAGE, prior to Western blotting as described above.

### Serum bactericidal assays

Serum bactericidal assays were performed four times using a modification of the previously described method^55^. Briefly, bacteria were grown to OD_600_= 2 in 5 ml of LB at 37°C and re-suspended in PBS to a final concentration of 10^7^ CFU/ml; 10 μl was then mixed with 90 μl of undiluted pooled healthy human serum at 37°C with shaking (180 rpm), and viable counts determined. Killing was confirmed to be due to the activity of complement by using 56°C heat-inactivated serum as a control.

### Chicken infection experiments

All work was conducted in accordance with United Kingdom legislation governing experimental animals under project license PPL 40/3652 and was approved by the University of Liverpool ethical review process prior to the award of the license. All birds were checked a minimum of twice daily to ensure their health and welfare. Birds were housed in accommodation meeting UK legislation requirements. 1-day old Lohmann Brown Layers were obtained from a commercial hatchery, separated into groups on arrival and given *ad libitum* access to water and a laboratory-grade vegetable protein-based pellet diet (SDS, Witham, UK). Chicks were housed at a temperature of 30°C. At 7 days of age chickens were inoculated by oral gavage with 10^8^ CFU of *S*. Typhimurium strains D23580, D23580 *pgtE*^P4/74^ or D23580 Δ*pgtE*. At 3 days post infection, 10 birds from each group were killed for post-mortem analysis. Samples from spleen, liver and the caecal contents were removed aseptically from each bird and diluted 1:5 (wt/vol.) in sterile phosphate-buffered saline (data from caecal content and liver not shown). Tissues were then homogenized in a Colworth 80 microstomacher (A.J. Seward & Co. Ltd, London, UK). Samples were serially diluted and dispensed onto Brilliant Green agar (Oxoid, Cambridge, UK) to quantify numbers of *Salmonella* as described previously^56^.

### Analysis of the conservation of the *pgtE* promoter SNP

259 genomes of *S.* Typhimurium ST313 isolates from Malawi and the UK^7^ were assembled using the A5 pipeline^57^ and Abacus^58^ unless reference quality genomes were available (all strains and accession numbers are given in Supplementary Table 2). Additionally 84 reference genomes were downloaded from NCBI that represent the known diversity of the *Salmonella* genus. Core genome SNPs were identified using the PanSeq package^59^ and a maximum likelihood phylogenetic tree was constructed from the concatenated SNP alignment using PhylML^60^. BLASTn was used to identify the genotype of the *pgtE* TSS −35 region nucleotide in all genomes (Supplementary Table 2).

## Acknowledgements

We are grateful to present and former members of the Hinton lab for helpful discussions, particularly Sathesh Srivasankaran and Aoife Colgan. We thank Paul Loughnane for his expert technical assistance, and Rob Kingsley for provision of strains. This work was supported by funding from a Wellcome Trust Senior Investigator award to JH (Grant 106914/Z/15/Z). DH was supported by the Wenner-Gren Foundation, Sweden. NW was supported by an Early Postdoc Mobility 543 fellowship from the Swiss National Science Foundation. RC was supported by an EU Marie Curie 544 International Incoming Fellowship (FP7-PEOPLE-2013-IIF, Project Reference 628450).

### Author Contributions

Conceived and designed experiments: DLH, CK, SVO and JCDH. Completed experiments and collected data: DLH, CK, SVO, LL, NW, TJW, PW and KH. Analysed and interpreted data: DLH, CK, SVO, RC, IRH, PW and JCDH. Wrote, critically revised or approved the final manuscript: DLH, CK, SVO, RC, LL, NW, TJW, IRH, PW, KH, NAF, MAG, JCDH.

## Supplementary data

**Supplementary Figure 1. Radial phylogeny illustrating the population structure of *S.* Typhimurium ST313 in the context of the *pgtE* promoter SNP.** The phylogenetic tree is reproduced with permission^7^. Red and blue coloured areas represent the presence of the T^D23580^ or C^4/74^ genotype respectively. Green coloured shading indicates the isolates belonging to the 313 sequence type.

**Supplementary Figure 1:** The *pgtE* promoter SNP in the context of *S.* Typhimurium ST313 population structure.

**Supplementary Table 1:** All TSS identified in D23580, and comparison to strain 4/74

**Supplementary Table 2:** SNPs and indels in the D23580 genome compared to the 4/74 genome

**Supplementary Table 3:** All SNPs within the −40 bp region of identified primary TSS in D23580 and PUV values for the respective TSS in D23580 and 4/74

**Supplementary Table 4:** Accession numbers, phylogenetic designation and *pgtE* TSS −40 sequence of all genomes sequences used in this study.

**Supplementary Table 5:** All bacterial strains and plasmids used in this study

**Supplementary Table 6:** All oligonucleotides used in this study

